# Unraveling the metabolic interactions of a *Dehalobacter*-containing anaerobic mixed culture for bioremediation

**DOI:** 10.64898/2026.05.05.723060

**Authors:** William T. Scott, Luz A. Puentes Jácome, Bart Nijsse, Jinsong Wang, Gerben R. Stouten, Jasper J. Koehorst, Hauke Smidt, Elizabeth A. Edwards, Peter J. Schaap, Robbert Kleerebezem

## Abstract

Organohalide-respiring bacteria (OHRB), such as *Dehalobacter*, play key roles in the bioremediation of anoxic environments contaminated with chlorinated aromatic compounds. These obligate anaerobes rely on syntrophic interactions to obtain essential resources—hydrogen, acetate, and corrinoid cofactors—from acetogens and fermenters. However, the metabolic interactions enabling complete reductive dehalogenation of compounds like 1,2,4-trichlorobenzene (1,2,4-TCB) to benzene remain incompletely understood. In this study, we asked: (1) What are the key microbial taxa and their functional roles within a *Dehalobacter*-containing anaerobic microbial community detoxifying chlorinated benzenes? (2) How do syntrophic interactions enable complete dehalogenation of 1,2,4-TCB to benzene under anaerobic conditions? (3) Can genome-resolved metagenomics and genome-scale metabolic modeling elucidate the metabolic dependencies supporting organohalide respiration in complex consortia? To address these questions, we cultivated microbial communities in batch reactors using methanol as electron donor and either 1,2,4-TCB or monochlorobenzene (MCB) as electron acceptor. In active MCB-fed cultures, benzene increased from 0 to 62.3µmol per bottle while MCB decreased from 88.3 to 22.0µmol per bottle over 120 days, with this pattern repeating across multiple substrate additions. Using genome-resolved metagenomics to identify dominant taxa and select 12 high-quality metagenome-assembled genomes (MAGs) for modeling, we reconstructed genome-scale metabolic models (GEMs) to identify candidate metabolic interactions and predict syntrophic dependencies that may support organohalide respiration in these consortia. Community flux sampling predicted that methanol, H_2_, acetate, and CO_2_ formed the dominant exchange backbone of the modeled community, while also indicating competition for shared electron donors between the two *Dehalobacter* populations. Model-guided minimal-community analysis further identified a narrow dechlorinating core in which all feasible minimal consortia retained a *Dehalobacter* member together with *Methanothrix*. These results provide a modeling-informed framework for hypothesis generation and future experimental validation of anaerobic consortia relevant to bioremediation.

## 1. Introduction

Chlorinated compounds historically used in industry and agriculture, such as chlorinated ethenes, ethanes, benzenes, and pesticides, are ubiquitous soil and groundwater pollutants (Aulenta et al., 2006). 1,2,4-Trichlorobenzene (1,2,4-TCB), a prominent example, is a chlorinated aromatic pollutant frequently detected at industrial sites, particularly in areas historically associated with chemical manufacturing, pesticide production, solvent use, and the disposal of chemical wastes (Zolezzi et al., 2005). Due to its chemical stability, resistance to degradation, and hydrophobic nature, 1,2,4-TCB readily accumulates in sediments and groundwater, posing substantial long-term risks to ecosystems and human health through toxicity, persistence, and bioaccumulation in food chains (Nelson et al., 2014). Conventional physicochemical remediation strategies, including chemical oxidation, thermal treatment, and physical extraction, have shown limited effectiveness against highly chlorinated aromatic pollutants due to their robust molecular structure and low solubility, highlighting the urgent need for alternative and cost-effective bioremediation strategies (Nelson et al., 2014).

Anaerobic biodegradation, specifically reductive dechlorination, has emerged as an effective and environmentally friendly method for addressing contamination by 1,2,4-TCB (Liu et al., 2025). This microbial process involves specialized organohalide-respiring bacteria (OHRB) that utilize chlorinated aromatic compounds as terminal electron acceptors in their energy metabolism, effectively removing chlorine atoms and transforming pollutants into progressively less toxic and more biodegradable intermediates (Alfán-Guzmán et al., 2017). Among OHRBs, the genus *Dehalobacter* has gained significant attention for its demonstrated ability to reductively dechlorinate 1,2,4-TCB to dichlorobenzene isomers (DCB), monochlorobenzene (MCB), and occasionally benzene, depending on strain-specific enzyme activities, available electron donors, and prevailing environmental conditions such as temperature, pH, and substrate availability (Nelson et al., 2014; Cao et al., 2025). For example, *Dehalobacter* sp. strain TeCB1 (isolated from 1,2,4,5-tetrachlorobenzene-contaminated ground-water) can grow by dechlorinating 1,2,4,5-TeCB to 1,2,4-TCB, then to 1,3- and 1,4-DCB (with 1,2-DCB and trace MCB as minor intermediates)(Alfán-Guzmán et al., 2017).

*Dehalobacter* spp. express distinctive reductive dehalogenase enzymes (RDases) that confer specific catalytic properties and substrate affinities toward chlorinated benzenes. Metaproteomic and metagenomic analyses have identified RDases, including the notable TcbA enzyme, which display unique evolutionary pathways compared to aromatic RDases from other OHRB genera, such as *Dehalococcoides* (Bulka et al., 2024; Li et al., 2025). More specifically, TcbA was found to be closely related (∼ 95% identity) to the PceA RDases of *Dehalobacter restrictus* and *Desulfitobacterium* spp., but only ∼ 19% similar to CbrA, the RDase in *Dehalococcoides mccartyi* strain CBDB1 that attacks chlorinated benzenes (Li et al., 2025). This distinct evolutionary lineage underscores *Dehalobacter*’s specialized metabolic adaptation to chlorinated aromatic pollutants, including unique catalytic mechanisms and substrate specificities (Alfán-Guzmán et al., 2017; Pérez-de Mora et al., 2018). Nonetheless,*Dehalobacter* alone may not fully dechlorinate 1,2,4-TCB, indicating the need for additional microbial partners to complete detoxification of the chlorinated intermediates formed (Qiao et al., 2019).

Anaerobic biodegradation mediated by *Dehalobacter* is highly dependent on the complexity and functionality of mixed microbial communities. These microbial consortia typically involve fermentative bacteria, which produce essential electron donors such as hydrogen and organic acids; methanogenic archaea, which play crucial roles in maintaining optimal redox conditions by scavenging excess hydrogen; and other bacteria responsible for the synthesis of essential cofactors, particularly corrinoids (vitamin B_12_), required by *Dehalobacter* for enzymatic function (Wang et al., 2017; Wen et al., 2020). One striking example of synergy is a mixed culture reported by Qiao et al. (2019) in which *Dehalogenimonas* and *Dehalobacter* populations worked in concert to completely dechlorinate 1,2,4-TCB. In this consortium, a *Dehalogenimonas* sp. was responsible for the initial dechlorination of 1,2,4-TCB to multiple DCB isomers (producing 1,2-DCB, 1,3-DCB, and 1,4-DCB), and could further dechlorinate 1,2-DCB and 1,3-DCB to monochlorobenzene. Other microbes in *Dehalobacter*-containing consortia provide essential support functions. Anaerobic fermenters produce the hydrogen or low-molecular-weight organics that serve as electron donors for reductive dechlorination. Such syntrophic interactions, metabolic interdependencies, and ecological cooperativity underscore the importance of community-based approaches rather than focusing solely on individual microbial isolates for successful bioremediation (Maphosa et al., 2010).

Constraint-based community modeling frameworks developed throughout the years provide foundational tools to simulate metabolic exchanges and cooperative behavior in microbial consortia, advancing the mechanistic study of *Dehalobacter*’s ecological role (Finley et al., 2010; Dukovski et al., 2021). Moreover, GEMs constructed for *Dehalobacter* strains offer computational platforms to predict growth conditions, interspecies interactions, and nutrient exchanges within microbial communities (Bulka et al., 2025). A recent study employed constraint-based metabolic modeling with a multi-omics approach to dissect a *Dehalobacter*-dominated enrichment culture degrading 1,2-dichloroethane and revealed how putative specialization and interspecies interactions drive stable dechlorination activity (Wang et al., 2019). More broadly, recent reviews emphasize the growing importance of systems biology approaches in elucidating the metabolism of anaerobes like *Dehalobacter*, in which integration of omics data with metabolic modeling enables predictive understanding of ecological function and bioremediation potential (Wang et al., 2023; Scott Jr et al., 2023). Together, these integrative studies provide a mechanistic basis for understanding *Dehalobacter*’s ecological function and support the rational design of bioremediation strategies in contaminated environments.

Despite significant advancements, knowledge gaps persist regarding ecological factors limiting *Dehalobacter*-driven biodegradation, resilience and stability of microbial consortia, and mechanisms underlying electron donor supply, nutrient cycling, and cofactor availability *in situ* (Saiyari et al., 2018). To bridge these gaps, the present study employed comprehensive metagenomic sequencing and genome-centric analyses on anaerobic enrichment cultures that dechlorinate 1,2,4-TCB to a mixture of DCBs and MCB. These cultures were derived from the laboratory culture KB-1 which dechlorinates chlorinated ethenes and ethanes. KB-1 was originally derived from a chlorinated compounds-contaminated environment. Previously published work demonstrated that the dechlorination of 1,2,4-TCB to DCBs and benzene occurred via organohalide respiration resulting in the growth of *Dehalobacter* (Puentes Jácome and Edwards, 2017). Similarly, MCB dechlorination to benzene was also found to occur through organohalide respiration, resulting in the growth of *Dehalobacter*(Puentes Jácome et al., 2021). High-quality MAGs were generated to investigate genomic diversity, metabolic versatility, and the overall functional potential within these microbial consortia.

Subsequently, GEMs were reconstructed and curated by incorporating experimentally supported pathways relevant to chlorinated benzene dehalogenation, and the resulting models were analyzed using pairwise and community-level constraint-based modeling. In particular, we applied an RHMC (Riemannian Hamiltonian Monte Carlo)-based flux-sampling framework to quantify pairwise ecological interactions and community metabolite exchange under pre-reduced defined mineral (PRDMM) medium constraints, following recent flux-sampling approaches for microbial communities (Gelbach et al., 2024; Klitgord and Segrè, 2010; Orth et al., 2010; Heirendt et al., 2019). By integrating these constraint-based modeling analyses with experimental data—including anaerobic cultivation, specialized DNA extraction, and high-throughput sequencing— this work provides a systems-level view of microbial interactions, metabolic exchange networks, and ecological dependencies that may govern anaerobic 1,2,4-TCB dechlorination and inform strategies for improved bioremediation.

## 2. Materials and methods

### 2.1. Culture harvesting and DNA extraction

Inside the anaerobic chamber, 3 × 50 mL were harted from each observation unit/enrichment culture (KB1_ CB_MeOH_2014, KB1_TCB_MeOH_2016, KB1_MCB_ MeOH_2017) and aliquoted into three 50 mL Falcon tubes. Falcon tubes were tightly sealed, and the caps were wrapped with anaerobic tape (for a pre-reduced defined mineral medium list, see Supplementary Data File S1). The samples were centrifuged outside the glove box at 6,100 × g for 30 min at room temperature. The Falcon tubes were then placed inside the glovebox and all but 10 mL of supernatant were discarded from each tube. The pellet was resuspended and combined into a single Falcon tube. The combined samples were centrifuged outside the glovebox at 6,100 × g for 10 min at room temperature. Subsequently, the Falcon tubes were placed inside the glove box and all but 1 mL of supernatant was discarded. The pellet was resuspended in 1 mL of culture and transferred to a 2 mL microtube with an O-ring screw cap. The samples were centrifuged at 10,000 rpm in a microcentrifuge for 10 min at room temperature. The pellet was resuspended in 250 µL of supernatant and stored at –80°C. The pellets were placed on ice for subsequent DNA extraction using the PowerSoil^®^ DNA extraction kit. The DNA extracts were submitted for whole metagenomics sequencing (Illumina paired-end sequencing) at the Beijing Genomics Institute. The DNA library was prepared using an internal preparation protocol with a 350 bp insert size and sequenced using the HiSeq 2500 sequencing platform (150 bp read length).

Figure 1 illustrates the enrichment culture setup history and the sequential dechlorination pathway observed in the (KB1_TCB_MeOH_2014, KB1_TCB_MeOH_2016, KB1_ MCB_MeOH_2017) cultures, showing the progression from 1,2,4-trichlorobenzene (1,2,4-TCB) to 1,3-dichlorobenzene (1,3-DCB), monochlorobenzene (MCB), and finally benzene, catalyzed by reductive dehalogenases (RDases).

**Figure 1:**
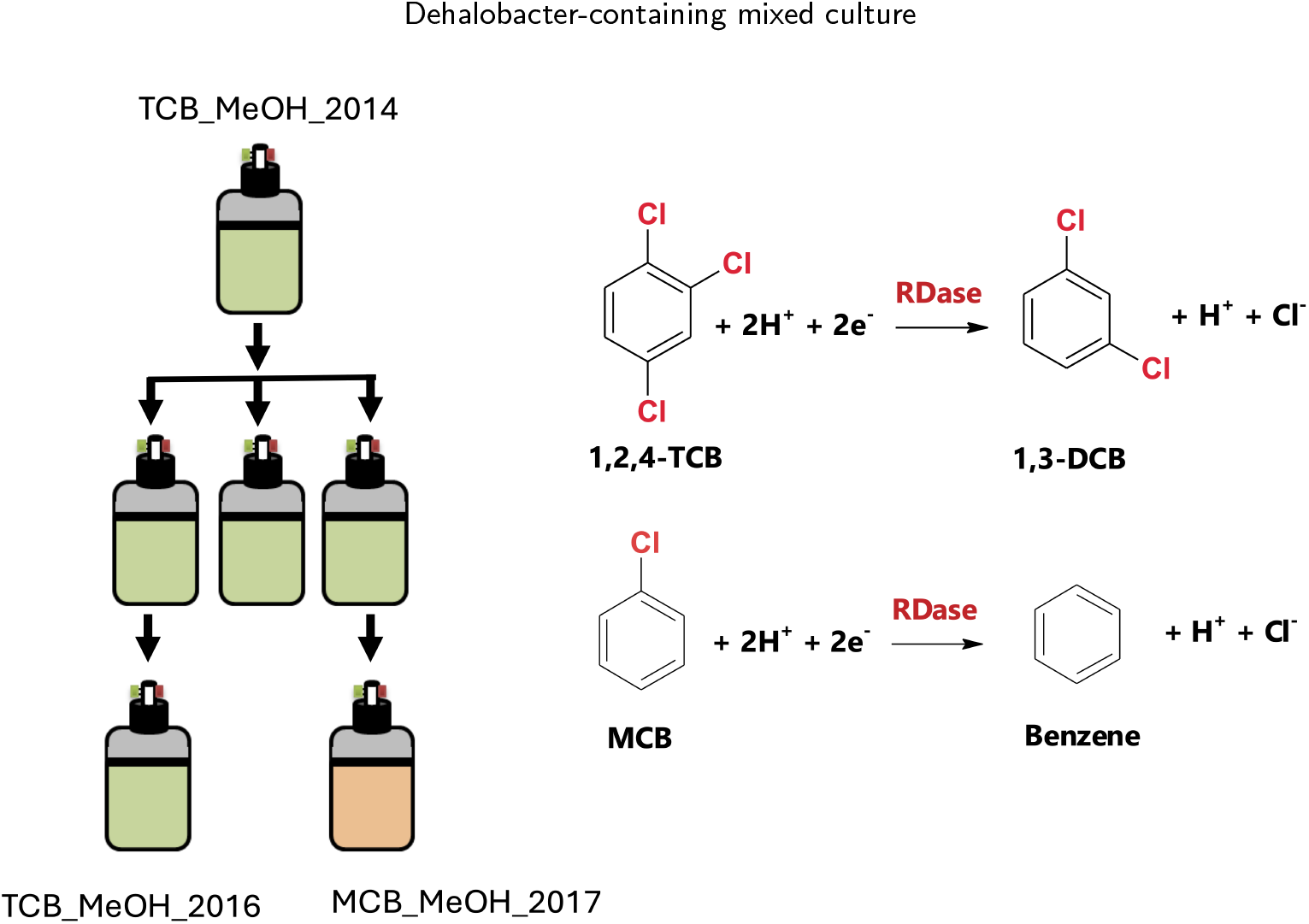
Enrichment culture history and reductive dechlorination framework. Enrichment culture history and critical reductive dechlorination reactions in KB-1 enrichment cultures catalyzed by reductive dehalogenase enzymes (RDases). Cultures are named following this convention: electron acceptor_electron donor_year established. TCB_MeOH_2014 is the parent culture of TCB_MeOH_2016 and MCB_MeoH_2017.

### 2.2. Genome-centric metagenomics

Metagenomic sequencing data from the three enrichment cultures (KB1_TCB_MeOH_2014, KB1_TCB_MeOH_2016, and KB1_MCB_MeOH_2017) were processed using a comprehensive genome-centric approach (Figure 2). The analysis pipeline was implemented following FAIR principles (Findable, Accessible, Interoperable, Reusable), with meta-data management and validation structured according to the FAIR Data Station framework (Nijsse et al., 2023). Raw sequencing reads were first assembled using meta-SPAdes with optimized k-mer settings to maximize contig length and minimize fragmentation of low-abundance genomes. Following assembly, reads from all three samples were mapped back to the assembled contigs using minimap2 with stringent alignment parameters (minimum mapping quality 30, minimum alignment length 50 bp) to accurately quantify contig coverage and abundance profiles across the different enrichment conditions.

**Figure 2:**
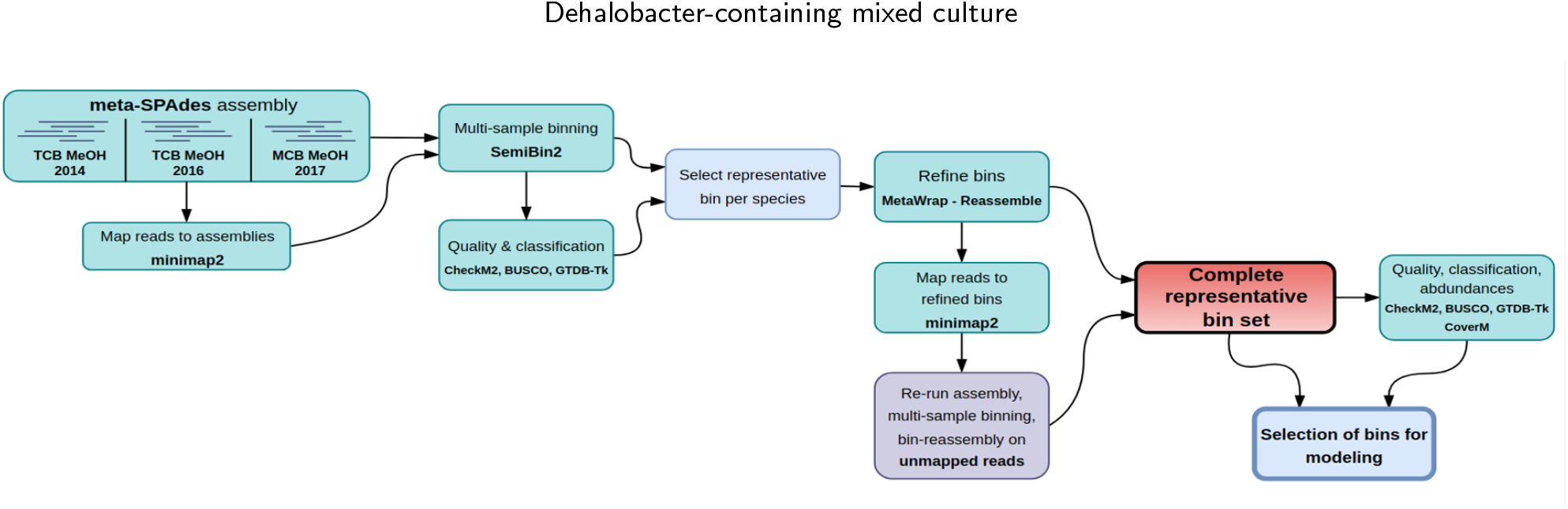
Metagenomic reconstruction and MAG selection workflow. Workflow for metagenomic assembly, binning, refinement, and selection of representative bins for modeling.

Multi-sample binning was performed using SemiBin2, which employs a semi-supervised deep learning approach that leverages both sequence composition features and differential coverage patterns to delineate individual genomes with high accuracy. The initial binning results were rigorously evaluated by CheckM2 (for contamination and completeness), BUSCO (for completeness of single-copy genes), and GTDB-Tk (for taxonomic classification). To improve genome quality and resolve strain-level variations, representative bins for each detected species were selected and refined using MetaWrap with the reassembly option, employing a minimum completion threshold of 80% and a maximum contamination threshold of 5%. For genomic regions not captured in initial binning, unmapped reads underwent additional assembly and binning iterations, increasing overall genome recovery by approximately 15%. The final set of representative bins was subjected to comprehensive quality assessment and abundance profiling using CoverM, with abundance values normalized to account for genome size and sequencing depth. Relative abundance was estimated by mapping quality-filtered reads back to the selected MAGs and calculating coverage-based abundance values normalized across samples.

### 2.3. Genome-scale metabolic reconstruction

High-quality MAGs (BUSCO score > 80% across the three samples) were selected for genome-scale reconstruction and *in silico* metabolic modeling. Detailed MAG properties are provided (Supplementary Data File S2).

Twelve GEMs were reconstructed from these MAGs using `gapseq` (Zimmermann et al., 2021). For each genome, the appropriate bacterial or archaeal reaction universe was used, and default gene-matching settings were applied to restrict pathway searches to the relevant taxonomic range. Where taxonomic assignment was sufficient, draft GEMs were generated with the corresponding biomass reaction type (gram-positive, gram-negative, or archaeal); otherwise, biomass composition was inferred automatically. Default `gapseq` bit-score thresholds (-l and -u) were used to distinguish reactions with or without sequence support.

Gap-filling was performed in pre-reduced defined mineral medium (PRDMM) to approximate the experimental conditions (Supplementary Data File S3), using a minimum bit score of 100 for inclusion of reactions in the core model. Reaction directionality was adjusted where appropriate to reflect experimental conditions, including elevated H_2_ availability. Futhermore, GEMs were constrained using a PRDMM derived from the laboratory recipe archived (Supplementary Data File S1) and translated into ModelSEED-compatible exchange bounds (Supplementary Data File S3). The formulation follows the bicarbonate-buffered, FeS-reduced mineral media used for KB-1 and related chlorobenzene-dechlorinating enrichments in the Edwards laboratory (Puentes Jácome and Edwards, 2017; Puentes Jácome, 2019). In principle, an extracellular concentration *C*_*i*_ could be related to a maximal uptake flux by 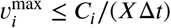, where *X* is biomass concentration and Δ*t* is the characteristic depletion time. Because these quantities are not uniquely defined for the enrichment-derived community simulations, the model medium was implemented as a semiquantitative constraint set: all experimentally supplied metabolites were made available for exchange, and upper uptake capacities were stratified by intended abundance class (major salts/buffer > primary substrates/gases > trace vitamins and cofactors). This preserves the experimental medium logic without imposing an arbitrary concentration-to-flux conversion.

Two GEMs, representing *Dehalobacter restrictus* and *Dehalobacter sp004343605*, were further curated to include known tri-, di-, and mono-chlorobenzene dehalogenation pathways. This curation was performed using `COBRApy` (Ebrahim et al., 2013) and `gapseq`, following a protocol similar to previous studies (Thiele and Palsson, 2010; Scott Jr et al., 2020, 2025). Detailed lists of added metabolites and reactions are provided in Supplementary Data Files S4 and S5. Basic properties of the final GEMs, including numbers of metabolites, reactions, genes, and compartments, are summarized in Table 1.

**Table 1.** Comparison of reaction, metabolite, and gene counts in GEMs reconstructed using GS (Gapseq) and CM (CarveMe).

| Model Name | GS Rxns | GS Mets | GS Genes | CM Rxns | CM Mets | CM Genes |
| --- | --- | --- | --- | --- | --- | --- |
| g_Methanothrix_s__ | 1437 | 1388 | 430 | 1049 | 786 | 491 |
| Methanobacterium_C_congolense | 1600 | 1553 | 429 | 1002 | 743 | 454 |
| Acetobacterium_sp003260995 | 1748 | 1513 | 558 | 1700 | 1095 | 901 |
| Dehalobacter_restrictus_cur | 1540 | 1389 | 422 | 1527 | 1000 | 654 |
| Dehalobacter_sp004343605_cur | 1466 | 1355 | 391 | 1446 | 970 | 610 |
| o_Bacteroidales_f_sp002418865 | 1649 | 1447 | 337 | 1603 | 1094 | 540 |
| o_Bacteroidales_g_LD21 | 2051 | 1681 | 536 | 2092 | 1364 | 855 |
| o_Bacteroidales_g_SR-FBR-E99 | 1924 | 1624 | 446 | 1996 | 1340 | 724 |
| Cloacimonadaceae_g_Cloacimonas | 1161 | 1086 | 242 | 1486 | 1054 | 401 |
| Rectinema_sp002441395 | 1696 | 1481 | 463 | 1834 | 1217 | 829 |
| Rectinema_sp015657395 | 1819 | 1594 | 472 | 2117 | 1401 | 919 |
| Rectinema_subterraneum | 1636 | 1374 | 436 | 1791 | 1155 | 800 |

### 2.4. Pairwise ecological interaction modeling

Pairwise ecological interactions were quantified using compartmentalized co-culture framework adapted from prior multi-species constraint-based approaches and the flux-sampling workflow of Gelbach *et al*. (Klitgord and Segrè, 2010; Gelbach et al., 2024; Orth et al., 2010). For each pair, the two GEMs were merged into a shared stoichiometric system with a common extracellular compartment and constrained to PRDMM using ModelSEED-compatible exchange mappings. Simulations were run in MATLAB with the COBRA Toolbox and Gurobi (Heirendt et al., 2019). Monoculture reference states were computed within each paired model by inactivating one partner while preserving the paired structure and constraints, yielding pair-specific monoculture growth distributions (Gelbach et al., 2024). After imposing a basal growth lower bound of 10% of the paired-model monoculture optimum for each organism, feasible flux states were sampled using constrained Riemannian Hamiltonian Monte Carlo (RHMC) (Gelbach et al., 2024; Heirendt et al., 2019). Ecological effects were classified using a 10% threshold on paired versus monoculture growth, and interactions were assigned from the joint sign pattern of the two responses; the final label was the most frequent outcome across quantile-matched sampled states (Gelbach et al., 2024; Klitgord and Segrè, 2010).

### 2.5. Community-level flux sampling

Community-level cross-feeding was inferred from random flux sampling of the 12-member *Dehalobacter*-containing community model under the PRDMM. The community model comprised organism-specific intracellular networks connected through a common extracellular compartment, such that secretion and uptake could be tracked across all taxa within a shared physicochemical environment (Orth et al., 2010; Klitgord and Segrè, 2010). Rather than relying on a single biomass-optimal solution, metabolite exchange was quantified across an ensemble of feasible steady-state flux distributions, following the rationale of Gelbach *et al*. that flux sampling better captures heterogeneity and submaximal-growth phenotypes in microbial communities (Gelbach et al., 2024). A metabolite was considered cross-fed when one organism secreted it to the extracellular compartment and another consumed it in the same sampled state.

For each donor–metabolite–receiver combination, exchange statistics were summarized across the sampled ensemble as mean flux, median flux and standard deviation. Exchange rates are reported as biomass-normalized model fluxes in mmol gDW^−1^ h^−1^, where gDW denotes grams dry-weight biomass. To focus on robust interactions, only exchanges with positive lower-bound support, defined conservatively as median – SD > 0, were retained for downstream network analysis and visualization. This procedure yielded reduced cross-feeding networks that resolved both the dominant catabolic exchange backbone and metabolite-specific subnetworks while preserving the variability inherent to the sampled feasible space (Gelbach et al., 2024).

### 2.6. Model-guided identification of minimal dechlorinating communities

Minimal dechlorinating communities were identified using an adapted `minMicrobiome` workflow (Raghu et al., 2024), which applies iterative taxon deletion and mixed-integer linear programming (MILP) to minimize community membership while retaining a target metabolic function. A 12-member community model built from ModelSEED-style SBML GEMs was constrained using PRDMM and evaluated under benzene or weighted dechlorination objectives. Simulations enforced uptake of 1,2,4-TCB, supplied methanol as the carbon and electron-donor substrate, and required retention of 80% of full-community growth and target-production capacity (`gr_frac` = 0.8; `scfa_frac` = 0.8) and 99% of the optimal growth solution space (`gr_opt_frac` = 0.99). Benzene was optimized either alone or jointly with chlorobenzene (1:0.5), and repeated MILP runs were used to identify alternative minimal solutions. Optimization was performed with Gurobi through the COBRA Toolbox.

## 3. Results

### 3.1. Observed reactor performance and study design

The biotransformation of chlorinated benzenes and the concurrent production of methane by different enrichment cultures over time are illustrated (Figure 3). The experimental setup for these enrichment cultures is detailed in Figure 1. Figure 3 comprises three panels, each showing the activity of a specific culture, with arrows indicating chlorinated benzene feeding events and magenta stars marking DNA sampling time points. Data points enclosed by circles represent calculated values rather than directly measured values.

**Figure 3:**
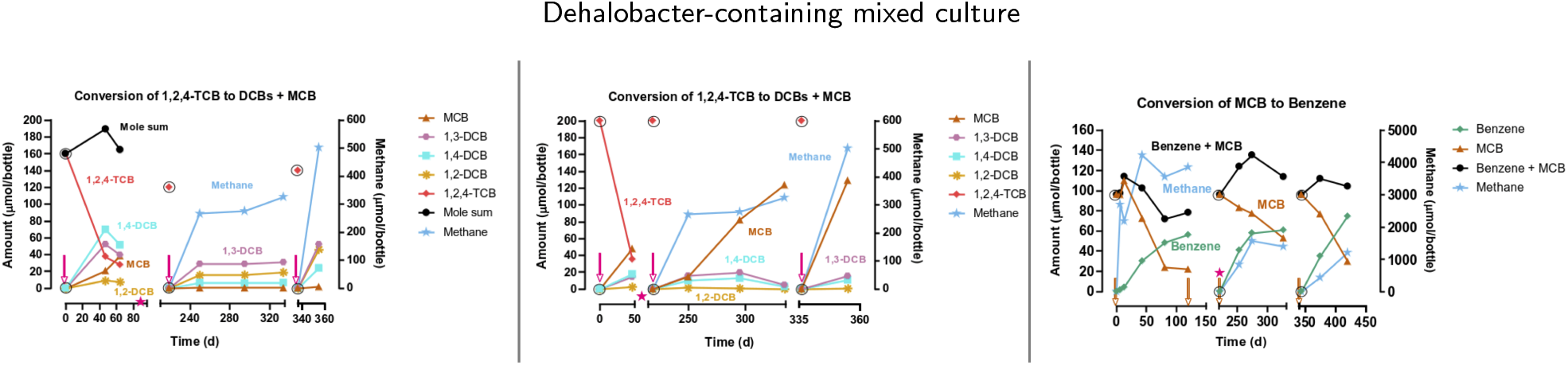
Biotransformation of chlorinated benzenes and methane production by enrichment cultures over time. **(Left, Panel 1)**: Conversion of 1,2,4-trichlorobenzene (1,2,4-TCB) to dichlorobenzenes (DCBs) and monochlorobenzene (MCB) by *KB1_TCB_MeOH_2014*. **(Center, Panel 2)**: Conversion of 1,2,4-TCB to DCBs and MCB by *KB1_TCB_MeOH_2016*. **(Right, Panel 3)**: Conversion of MCB to benzene by *KB1_MCB_MeOH_2017*. In all panels, arrows indicate chlorinated benzene feeding events, which were preceded by sparging with N_2_/CO_2_ to remove volatiles. Magenta stars indicate timepoints for DNA sampling (Day 92 in Panels 1–2; Day 221 in Panel 3), with feeding occurring after sampling in Panel 3. Data points enclosed by circles represent calculated rather than directly measured values. 1,2,4-TCB was measured only during the first cycle in Panels 1 and Methane production was monitored throughout, while total chlorinated compound recovery is shown as “Mole sum” (Panels 1 and 3) or inferred from product profiles.

As shown in Figure 3, Panels 1 and 2 depict the conversion of 1,2,4-TCB by enrichment cultures *KB1_TCB_Me OH_2014* and *KB1_TCB_MeOH_2016*, respectively. In both panels, 1,2,4-TCB concentrations (measured only during the first cycle) decreased rapidly after initial feeding events, leading to the formation of various DCBs, including 1,3-DCB, 1,4-DCB and 1,2-DCB, as well as MCB. The “Mole sum” in Panel 1 tracks the total chlorinated compounds recovered, while in Panel 2, transformation is inferred from product profiles. Methane production, shown in light blue, increased over time in both cultures, particularly after feeding events. DNA sampling occurred on Day 92 in both panels.

Panel 3 of Figure 3 shows the *KB1_MCB_MeOH_2017* enrichment culture, in which MCB decreased over successive feeding cycles while benzene increased. Following MCB:MeOH additions around days 120, 220, and 340, MCB declined and benzene accumulated, producing a consistent inverse relationship between the two compounds. From the start of the incubation, MCB decreased from approximately 100 µmol bottle^−1^ to about 22 µmol bottle^−1^, whereas benzene increased from near 0 to approximately 60 µmol bottle^−1^ by day 140. This pattern was observed again after subsequent feedings. The “Benzene + MCB” trace shows the combined total of these compounds. Methane also increased over time in this culture. DNA was sampled on day 221, and feeding was performed after sampling.

As shown in Figure 3, methane production increased in all active cultures during the incubation period. In the methanol-fed enrichments, increases in methane were observed following feeding events and during periods when chlorinated benzenes were also being transformed. This temporal pattern is consistent with the raw data provided in Supplementary Data File S6 and S7 (`TCB_MeOH_2014` and `MCB_MeOH` Excel spreadsheets).

### 3.2. Taxonomic and functional composition of the microbiome

Metagenomic analysis of the three methanol-fed enrichment cultures recovered high-quality MAGs spanning a taxonomically diverse, but functionally coherent anaerobic community centered on organohalide respiration (Table 2; Figure 4). Recovered MAGs showed BUSCO completeness values of 81.5–100% and low CheckM2 contamination (0.16–2.64%). Across the enrichments, the community was consistently structured around *Dehalobacter*, methanogenic archaea, and a set of fermentative or acetogenic bacteria.

**Table 2.** Assembly statistics and estimated relative abundances of MAGs recovered from methanol-fed anaerobic enrichment cultures. **Largest** refers to the size (bp) of the largest contig; **TCB 2014, TCB 2016**, and **MCB 2017** indicate the relative abundance (%) of each MAG in the three corresponding enrichment communities. Completeness and contamination were estimated using CheckM2. BUSCO taxonomic lineage and completeness scores (C) are also reported.

| Short | Contigs | Size | Largest | N50 | GC | TCB 2014 | TCB 2016 | MCB 2017 | BUSCO_Tax | BUSCO_score | CheckM2 Comp. | CheckM2 Contam. |
| --- | --- | --- | --- | --- | --- | --- | --- | --- | --- | --- | --- | --- |
| *Methanotrix* | 51 | 2,814,811 | 382,134 | 75,962 | 51.7 | 2.44 | 0.02 | 0.06 | methanomicrobia_odb10 | C:95.1% | 99.88 | 0.87 |
| *Methanobacterium_C congolense* | 6 | 2,397,603 | 1,069,109 | 845,458 | 38.5 | 3.89 | 2.01 | 0.31 | methanobacteria_odb10 | C:98.9% | 99.95 | 0.50 |
| *Acetobacterium sp003260995* | 43 | 3,802,546 | 363,528 | 187,074 | 42.5 | 2.66 | 0.05 | 0.03 | clostridiales_odb10 | C:98.1% | 99.97 | 1.35 |
| *Dehalobacter restrictus* | 41 | 3,185,342 | 336,294 | 106,851 | 44.4 | 36.73 | 1.62 | 0.09 | clostridiales_odb10 | C:98.1% | 99.99 | 1.77 |
| *Dehalobacter sp004343605* | 10 | 2,726,007 | 923,758 | 558,892 | 43.9 | 0.34 | 45.23 | 2.74 | clostridiales_odb10 | C:99.3% | 98.52 | 0.75 |
| *o_Bacteroidales; TTA-H9* | 17 | 2,823,461 | 472,940 | 206,935 | 37.8 | 1.27 | 2.95 | 5.33 | bacteroidetes_odb10 | C:96.2% | 98.90 | 2.64 |
| *o_Bacteroidales; LD21* | 23 | 4,968,462 | 644,554 | 517,608 | 41.7 | 0.01 | 0.01 | 1.29 | bacteroidia_odb10 | C:96.0% | 98.52 | 0.75 |
| *o_Bacteroidales; SR-FBR-E99* | 26 | 3,900,019 | 409,054 | 198,237 | 49.7 | 0.08 | 0.51 | 10.52 | bacteroidia_odb10 | C:96.6% | 99.05 | 0.49 |
| *Cloacimonas* | 11 | 2,121,983 | 757,500 | 238,605 | 39.1 | 2.51 | 2.92 | 13.71 | bacteria_odb10 | C:81.5% | 93.47 | 0.61 |
| *Rectinema sp002441395* | 13 | 2,857,512 | 498,091 | 272,893 | 53.1 | 0.63 | 4.25 | 0.42 | spirochaetales_odb10 | C:83.8% | 97.56 | 2.17 |
| *Rectinema sp015657395* | 40 | 3,099,071 | 768,770 | 560,874 | 56.2 | 7.44 | 9.73 | 0.63 | spirochaetales_odb10 | C:83.8% | 95.34 | 1.44 |
| *Rectinema subterraneum* | 9 | 2,636,913 | 291,911 | 102,325 | 52.3 | 8.10 | 3.04 | 2.16 | spirochaetales_odb10 | C:82.9% | 99.06 | 1.30 |

**Figure 4:**
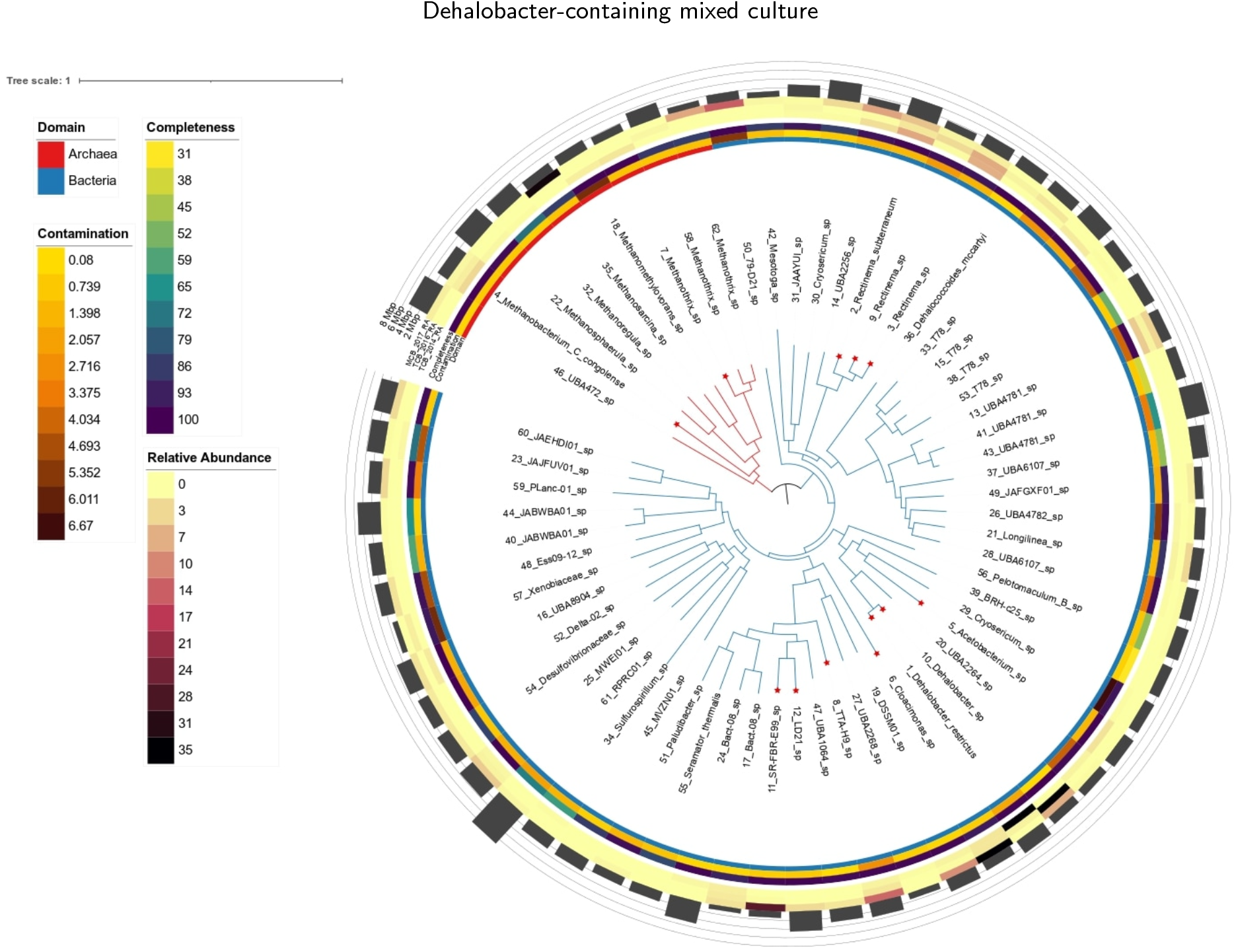
Phylogenomic relationships and genome quality patterns among recovered MAGs. Phylogenomic tree of all recovered MAGs from the three methanol-fed enrichment cultures. Branch colors indicate domain affliation (red = Archaea, blue = Bacteria). Concentric rings represent MAG completeness, contamination, and relative abundance across samples. Red asterisks mark the 12 MAGs selected for downstream genome-scale metabolic modeling, based on their taxonomic prominence, abundance, and high genome quality.

Two *Dehalobacter* populations dominated the dechlorinating guild and shifted across enrichments. *Dehalobacter restrictus* (Bin TCB_2014_SB_10_SELCT*) was most abundant in the 2014 TCB-fed culture (35.37%) but declined markedly thereafter, whereas *Dehalobacter* sp004343605 (Bin TCB_2016_SB_6_SELCT*) became dominant in the 2016 culture (46.2%) and remained detectable in 2017 (2.57%). Phylogenetic placement supported their separation within the genus (Figure 4), and these metagenomic results are consistent with earlier 16S rRNA gene analyses that detected two distinct *Dehalobacter*-affiliated sequence types (Puentes Jácome and Edwards, 2017). Methanogenic archaea formed a secondary guild: *Methanothrix* sp002498745 was more abundant in the 2014 culture and nearly absent from the later enrichments, whereas *Methanobacterium_C congolense* remained detectable in the TCB-fed cultures before declining in the MCB-fed culture.

Several fermentative and acetogenic taxa also varied across enrichments. *Acetobacterium* sp003260995 was present at moderate abundance in 2014 (2.58%) but diminished thereafter, whereas *Cloacimonas* increased from 2.43% to 12.8% over time. Three phylogenetically distinct *Rectinema* populations were prevalent in the TCB-fed cultures. Based on abundance, phylogenetic representation, and genome quality, 12 MAGs were selected for downstream metabolic modeling (completeness >95%, contamination <3%), including dominant *Dehalobacter, Methanobacterium, Methanothrix, Acetobacterium, Cloacimonas*, and *Rectinema* lineages (Figure 4).

### 3.3. Genome-guided metabolic reconstruction

To evaluate the metabolic potential of dominant taxa in methanol-fed mixed anaerobic dechlorinating communities, GEMs were reconstructed for twelve high-quality MAGs. Automated reconstruction was performed using both the Gapseq (GS) and CarveMe (CM) pipelines, each leveraging genome annotations and curated metabolic reaction databases. The resulting models varied in size and complexity, with GS-based models comprising between 1,161 and 2,051 reactions, 1,086 to 1,681 metabolites, and 242 to 558 annotated genes (Table 1). CM GEMs generally included fewer genes, though some exhibited a higher number of total reactions, particularly for certain Bacteroidales and *Rectinema* strains.

All GEMs were subjected to a comprehensive qual-assessment, including stoichiometric consistency, mass and charge balance, flux feasibility in minimal medium, and MEMOTE scoring (Table 3). Although CM GEMs achieved slightly higher MEMOTE scores in isolated cases, GS GEMs consistently demonstrated superior biochemical soundness. In particular, all GS GEMs exhibited complete stoichiometric consistency (100%) and near perfect mass and charge balance (≥99.8%), whereas multiple CM GEMs showed inconsistencies, including three with 0% stoichiometric consistency. Additionally, GS GEMs exhibited fewer reactions with unbounded flux under default medium conditions, suggesting improved thermodynamic feasibility. Moreover, GS GEMs offered more comprehensive gene-reaction associations, particularly for taxa involved in syntrophic acetate and hydrogen metabolism. Based on these cumulative advantages, including greater structural consistency, biochemical completeness, and environmental relevance, the GS-derived GEMs were selected for downstream community modeling and simulation analyses.

**Table 3.**
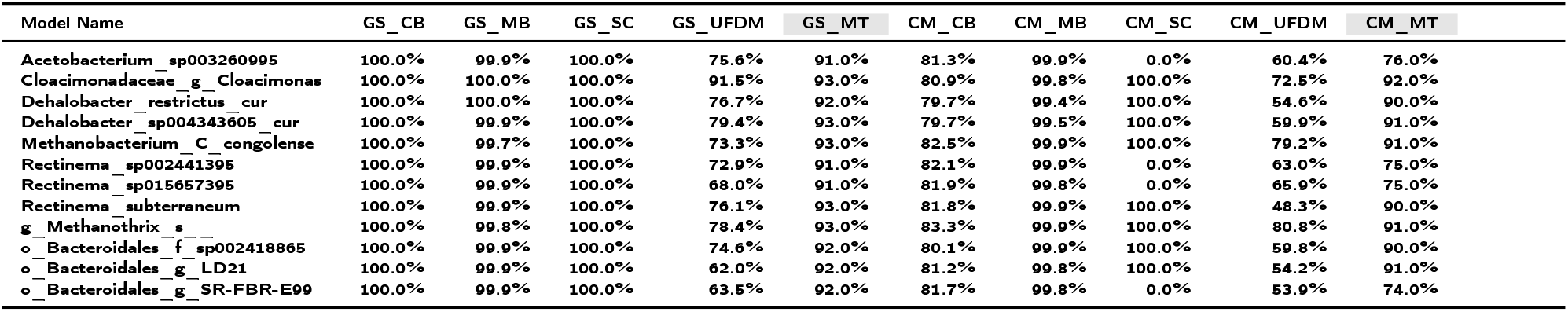
Compact summary of GEMs reconstructed using **GS** (Gapseq) and **CM** (CarveMe) pipelines. Metrics include **MT** (MEMOTE score), **SC** (Stoichiometric Consistency), **MB** (Mass Balance), **CB** (Charge Balance), and **UFDM** (Unbounded Flux in Default Medium). **Note: MEMOTE score excludes the Annotation - Genes subscore from the total score estimation.** MEMOTE scores are highlighted in gray.

To further characterize metabolic similarities across the reconstructed models, pairwise Jaccard indices were computed based on shared reaction content. The resulting clustered heatmap (Figure 5) revealed patterns of both functional similarity and divergence across taxa. As expected, the two *Dehalobacter* strains exhibited very high similarity (Jaccard index ε0.9), consistent with their phylogenetic proximity and shared reductive dehalogenation capabilities. Methanogens such as *Methanothrix* and *Methanobacterium* formed distinct clusters, reflecting their specialized roles in acetoclastic and hydrogenotrophic methanogenesis, respectively.

**Figure 5:**
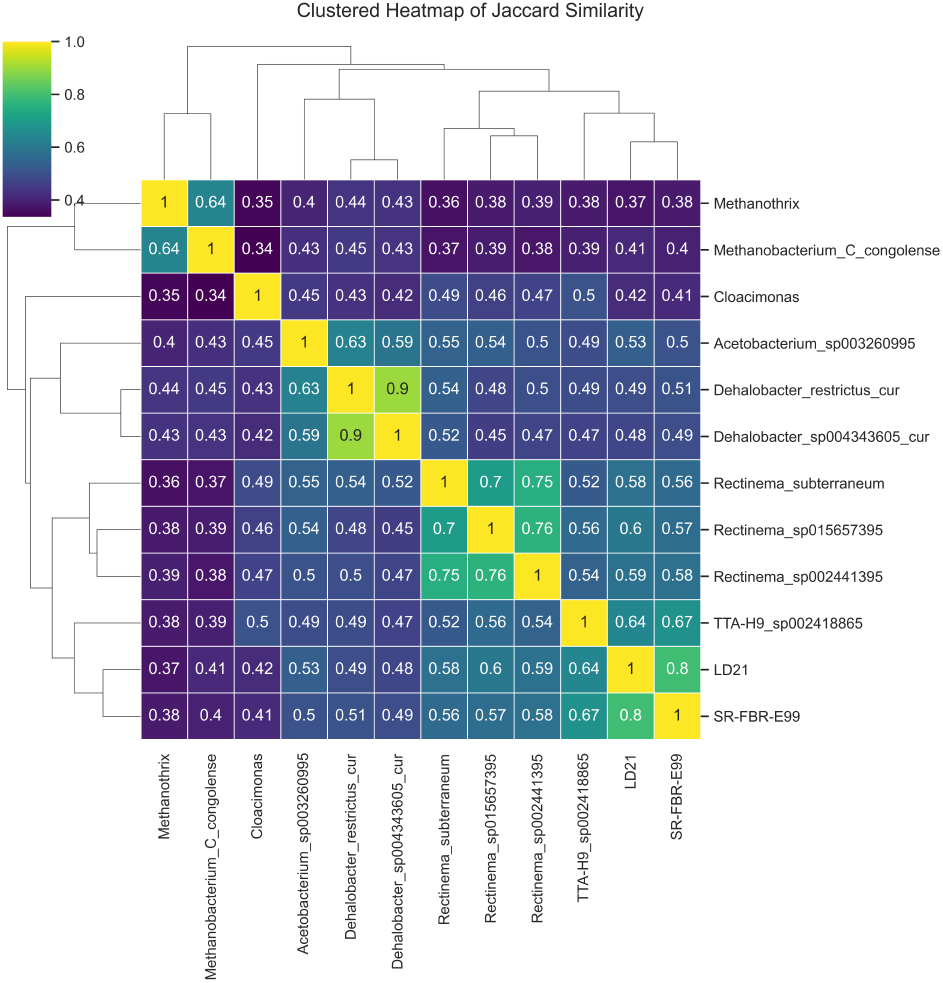
Pairwise similarity of metabolic reaction repertoires among reconstructed GEMs. Clustered heatmap of Jaccard similarity indices comparing metabolic reaction content among the 12 reconstructed GEMs. Values closer to 1.0 indicate higher overlap in reaction presence.

Interestingly, the three *Rectinema* strains clustered together, but displayed a broader range of Jaccard similarity values (0.70–0.76) than typically observed among genomes from the same genus. This modest level of overlap suggests underlying metabolic differentiation, possibly due to strain-specific adaptation. Closer inspection of their metabolic networks revealed differences in carbohydrate utilization, amino acid biosynthesis pathways, and presence or absence of lactate and ethanol fermentation routes. These differences do not appear to stem from genome contamination, as the MAGs exhibited high completeness and low contamination scores (Supplementary Data File S2). Altogether, these patterns highlight both conserved and divergent metabolic traits that contribute to a functionally specialized and cooperative microbial community tailored for anaerobic methanol degradation.

### 3.4. Random flux sampling reveals a catabolic exchange backbone

Random flux sampling of the 12-member community, conservatively filtered to retain only exchanges with positive lower-bound support (median – SD > 0), resolved a robust metabolite-restricted network comprising 377 significant directed interactions. Although broader than the earlier summary network, exchange activity remained strongly concentrated in a small catabolic core: H_2_, MeOH, EtOH, H^+^, acetate, and CO_2_ together accounted for 90.7% of the summed median interaction strength (1391.9 of 1534.4 mmol gDW^−1^ h^−1^) and 74.3% of all significant edges (280 of 377). H_2_ was the largest hub (326.3 mmol gDW^−1^ h^−1^; 21.3% of total median strength), followed by MeOH (302.3; 19.7%) and EtOH (279.3; 18.2%), with H^+^, acetate, and CO_2_ forming a second tier (Figure 6). These results indicate that feasible community states are repeatedly organized around a small set of shared catabolic metabolites that dominate community-level exchange flux.

**Figure 6:**
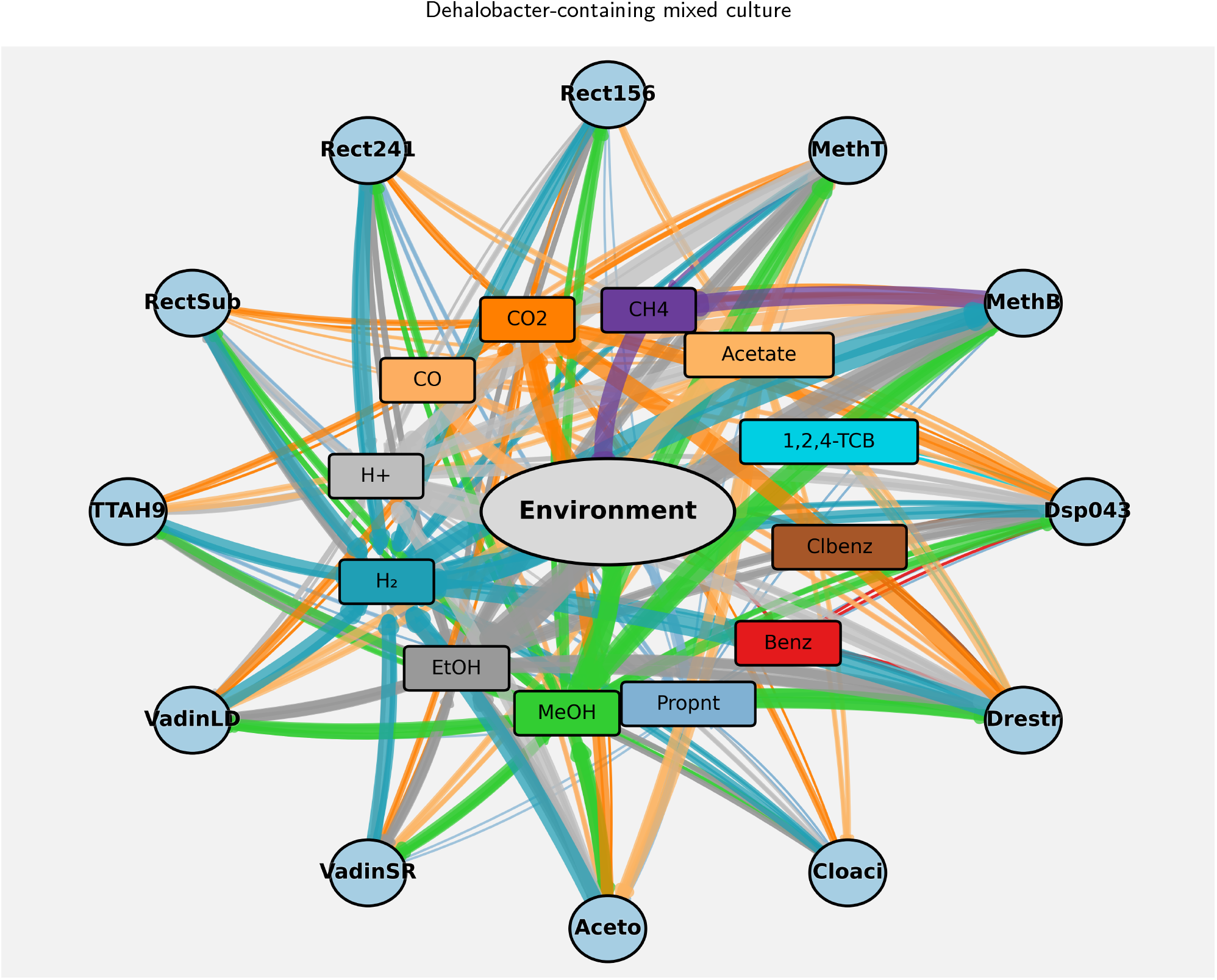
Significant metabolite cross-feeding network from community-level random flux sampling. Cross-feeding interactions in the 12-member dechlorinating community were inferred from community-level random flux sampling and filtered to retain only statistically robust exchanges with positive lower-bound support (median – SD > 0). Circular nodes represent microbial populations, the central ellipse denotes the shared extracellular environment, and square nodes represent metabolites involved in significant exchange. Directed edges indicate metabolite release to, or uptake from, the extracellular compartment, and edge thickness is proportional to exchange magnitude. The species abbreviations denote the modeled community members as follows: MethB, *Methanobacterium C. congolense*; MethT, *Methanothrix* sp.; Aceto, *Acetobacterium* sp003260995; Drestr, *Dehalobacter restrictus*; Dsp043, *Dehalobacter* sp004343605; Cloaci, *Cloacimonas*; Rect241, *Rectinema* sp002441395; Rect156, *Rectinema* sp015657395; RectSub, *Rectinema subterraneum*; TTAH9, o__Bacteroidales; f__sp002418865 (TTA-H9 lineage); VadinLD, o__Bacteroidales; g__LD21; and VadinSR, o__Bacteroidales; g__SR-FBR-E99. Metabolite abbreviations are MeOH, methanol; EtOH, ethanol; H_2_, hydrogen; H^+^, proton; CO_2_, carbon dioxide; CH_4_, methane; 1,2,4-TCB, 1,2,4-trichlorobenzene; Clbenz, chlorobenzene; Benz, benzene; and Propnt, propionate.

This backbone was also taxonomically structured. *Methanoba terium C. congolense* was the dominant H_2_ sink (median import 26.35 mmol gDW^−1^ h^−1^), whereas *Acetobacterium* sp003260995 was the strongest H_2_ source (median export 20.55 mmol gDW^−1^ h^−1^). *Methanothrix* was the clearest acetate sink (median import 3.93 mmol gDW^−1^ h^−1^), while also showing strong MeOH uptake (9.90 mmol gDW^−1^ h^−1^) and the largest H^+^ export signal (28.50 mmol gDW^−1^ h^−1^). CH_4_ production was assigned mainly to *Methanobacterium* (median 24.83 mmol gDW^−1^ h^−1^), with a smaller contribution from *Methanothrix* (2.01 mmol gDW^−1^ h^−1^). Taken together, these patterns suggest that dechlorination is embedded within a cooperative metabolic network in which acetogens, fermenters, and methanogens help redistribute electron donors and fermentation products, thereby sustaining the physicochemical conditions and metabolite supply required for continued organohalide respiration by *Dehalobacter*.

By contrast, the chlorinated aromatic module remained quantitatively minor but topologically specialized. 1,2,4-TCB, chlorobenzene, and benzene collectively contributed only 16.3 mmol gDW^−1^ h^−1^ of summed median interaction strength, or about 1.1% of the network total, and each was represented by only 10 significant edges. These exchanges were restricted almost entirely to the two *Dehalobacter* populations, consistent with a localized organohalide-respiring niche embedded within a much larger catabolic scaffold. Within this module, *Dehalobacter* sp004343605 showed the strongest association with 1,2,4-TCB uptake (median 0.518 mmol gDW^−1^ h^−1^), whereas *D. restrictus* showed the larger CO_2_ export signal (14.51 mmol gDW^−1^ h^−1^) and substantial acetate export (5.76 mmol gDW^−1^ h^−1^). Overall, the model supports a division of labor in which chlorinated benzene transformation is carried out by a specialized dechlorinating guild, but depends on metabolic support from the surrounding 12-species community.

### 3.5. Model-guided identification of a minimal dechlorinating core

To explore how community-scale metabolic models could be reduced while retaining dechlorination function, we adapted *minMicrobiome* to identify the smallest consortia capable of sustaining TCB transformation in PRDMM with methanol as the supplied carbon and electron-donor substrate. All simulations enforced uptake of both TCB and methanol and required retention of 80% of full-community growth and target-production capacity (`gr_frac = 0.8`, `target_frac = 0.8`, `gr_opt_frac = 0.99`). Under a benzene objective, seven minimal communities were recovered, all of which contained a *Dehalobacter* population together with *Methanothrix* (Figure 7). The smallest feasible solution was a two-member *D. restrictus*–*Methanothrix* consortium, although most solutions contained three members, and predicted target flux remained unchanged at 0.100 across all benzene solutions.

**Figure 7:**
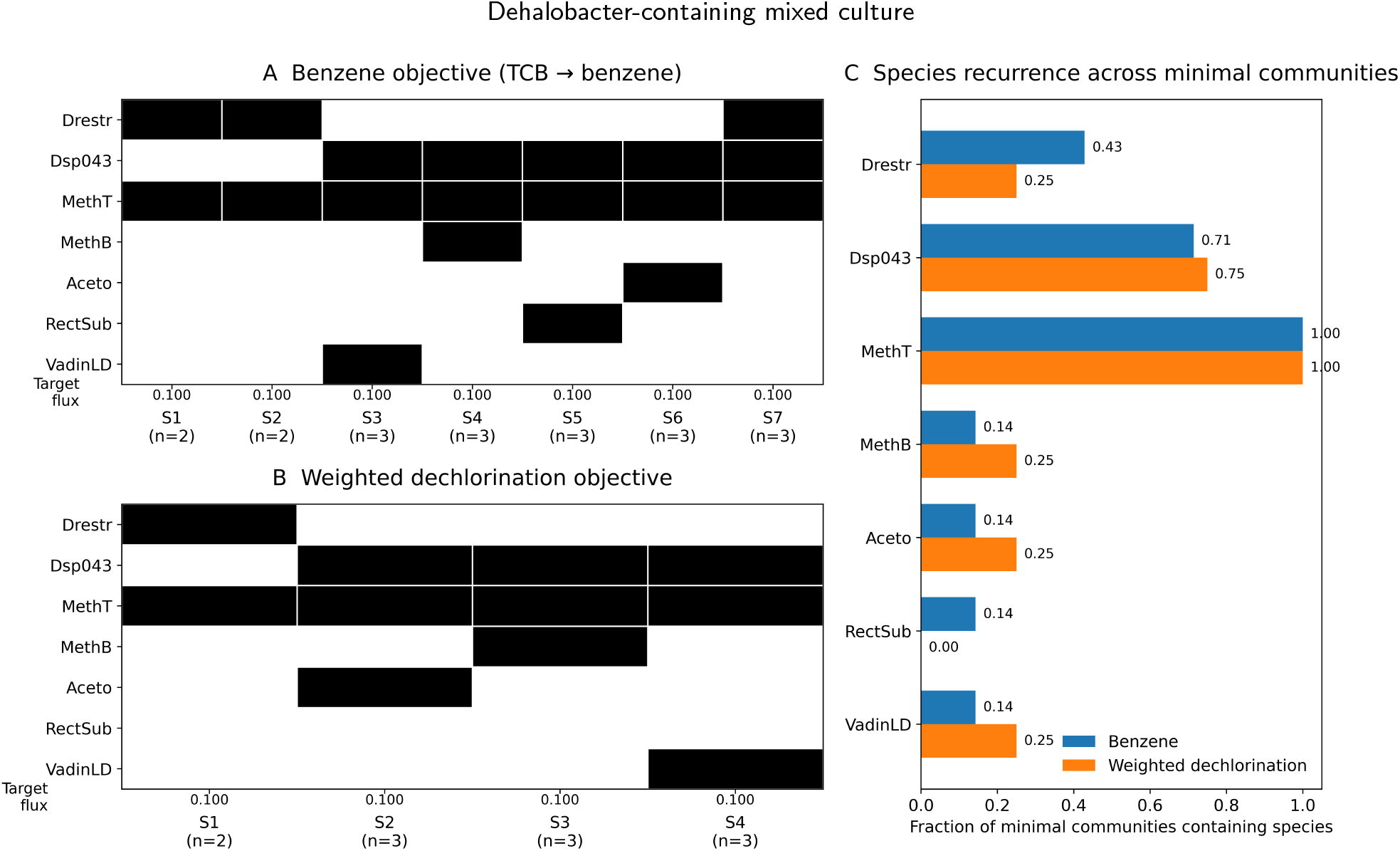
Model-guided minimal communities predicted for methanol-fed TCB dechlorination. Filled cells indicate membership in each minimal microbiome recovered by the adapted *minMicrobiome* workflow. All reported solutions were constrained to take up TCB and methanol. Across both the benzene and weighted-dechlorination objectives, every feasible minimal community retained a *Dehalobacter* member together with *Methanothrix*, whereas the smallest solution was the two-member consortium *D. restrictus* + *Methanothrix*. Three-member alternatives replaced the third taxon without materially changing the predicted target flux, indicating functional redundancy among auxiliary partners under the simulated medium.

A weighted dechlorination objective favoring benzene over chlorobenzene (1:0.5) produced four minimal com-m*c-*unities with a similar architecture (Figure 7). In these solutions, *Methanothrix* was retained in 100% of cases, and a *Dehalobacter* member was also retained in 100% of solutions when both *Dehalobacter* populations were considered together (Figure 7C), whereas *Methanobacterium, Acetobacterium, Rectinema*, and the VadinHA17-affiliated population each occurred in only 14–25% of solutions. Collectively, these results define a set of model-minimal candidate consortia for methanol-fed TCB transformation under the simulated medium and objective function, rather than an experimentally confirmed minimal biological community.

## Discussion

### Microbial drivers of dechlorination

This study examined the structure, functional roles, and metabolic dependencies of anaerobic enrichment cultures capable of dechlorinating 1,2,4-TCB and MCB to benzene with methanol as the primary substrate. By integrating genome-centric metagenomics with genome-scale metabolic modeling, we investigated how interactions among dechlorinators, methanogens, and fermentative or acetogenic community members may support chlorobenzene detoxification. Overall, the results indicate that effective dechlorination is embedded within a broader community context shaped by electron-donor transfer, metabolite exchange, and cofactor dependence, rather than by the dechlorinating populations alone.

Our investigation yielded several key findings. Firstly, we two confirmed the stable coexistence and central role of distinct *Dehalobacter* populations, closely related to *D. restrictus* and *Dehalobacter* sp004343605, as the primary agents driving the reductive dechlorination process from 1,2,4-TCB towards benzene. Secondly, we identified a core syntrophic network essential for supporting these *Dehalobacter* spp. This network comprises specific functional guilds, including hydrogen-producing fermenters (e.g., *Bacteroidales, Cloacimonas, Rectinema*), an acetogen capable of utilizing methanol derivatives (*Acetobacterium*), and methanogens responsible for consuming key intermediates like H_2_ and acetate (*Methanothrix* and *Methanobacterium*). Thirdly, our community metabolic modeling provided crucial insights into the functional dynamics, predicting essential nutrient requirements (notably cobalt for RDases), quantifying potential metabolic contributions of different members, and mapping the critical exchange of metabolites, particularly H_2_ and acetate, which structure the communityś syntrophic relationships. These findings directly address our research objective by providing a detailed, systems-level view of the microbial players and metabolic crossfeeding governing chlorobenzene dechlorination in this methanol-fed enrichment.

### 4.2. Coexistence and putative specialization of *Dehalobacter* spp

Our integrated metagenomic and modeling analyses revealed the coexistence of two distinct *Dehalobacter* populations, closely related to *Dehalobacter restrictus* and *Dehalobacter* sp004343605, within enrichments dechlorinating 1,2,4-TCB and MCB. This is notable because *Dehalobacter* species are generally considered metabolically specialized organohalide respirers with limited catabolic breadth (Wang et al., 2017; Bulka et al., 2024). The presence of two *Dehalobacter* lineages within the same enrichment suggests that chlorobenzene dechlorination was carried out by a mixed dechlorinating guild rather than a single dominant population. GEMs reconstructed from both MAGs supported their potential to use chlorinated benzenes as electron acceptors and H_2_ as the immediate electron donor, consistent with their central role in the observed dechlorination phenotype (Nelson et al., 2014).

Although both MAGs encoded dechlorination potential, their distinct *rdh* complements suggest non-identical functional roles. One encoded a characterized TcbA-like RDase associated with 1,2,4-TCB dechlorination, whereas the other harbored a candidate RDase potentially involved in MCB dechlorination, although this function remains unconfirmed. Different *Dehalobacter* strains are known to vary in chlorobenzene substrate range and dechlorination efficiency (Nelson et al., 2014; Alfán-Guzmán et al., 2017), and such differences may have contributed to the observed shift in dominance between the two populations. At the same time, pairwise interaction modeling predicted competitive or amensalistic interactions involving *Dehalobacter*, consistent with competition for limiting resources such as H_2_ or trace nutrients. In this context, coexistence does not necessarily imply efficient cooperation; rather, both populations may persist and dechlorinate under electron-donor-limited conditions in which H_2_ is shared among multiple community members, including methanogens, potentially constraining net dechlorination rates.

The basis for coexistence is therefore likely multifactorial and not fully captured by the current models. In addition to differences in *rdh* content, variation in RDase expression, corrinoid dependence, growth yield, and microscale spatial structure may influence which population dominates under a given enrichment history. Related studies have shown that multiple organohalide-respiring populations can coexist when different chlorinated intermediates or dechlorination steps are available within the same system (Qiao et al., 2019; Bulka et al., 2025). In the present study, however, the specific physiological basis of coexistence remains unresolved and will require direct validation, for example through metatranscriptomic profiling or targeted physiological assays.

### 4.3. Syntrophic network supporting dechlorination

Successful organohalide respiration by specialists like *Dehalobacter* typically depends on metabolic crossfeeding within a microbial community (Maphosa et al., 2010; Liu et al., 2025). In our community-level metabolic models, the cross-feeding network (Figure 6) was centered on methanol, H_2_, acetate, and CO_2_, indicating that dechlorination is embedded within broader community metabolism rather than driven by the dechlorinators alone. Key partners included methanogens (*Methanothrix, Methanobacterium*), an acetogen (*Acetobacterium*), and fermentative bacteria from lineages such as *Bacteroidales, Cloacimonas*, and *Rectinema*. H_2_ and acetate emerged as central intermediates, consistent with H_2_ serving as the immediate electron donor for *Dehalobacter* reductive dechlorination, while acetate contributes to broader carbon flow through the consortium (Wang et al., 2019; Qiao et al., 2019; Xu et al., 2024). The data further support a pathway-level division of labor in methanol utilization: *Acetobacterium* was the clearest candidate for primary methanol conversion, including the potential conversion of methanol and CO_2_ to acetate via the Wood–Ljungdahl pathway, whereas *Methanobacterium* was more plausibly associated with H_2_/CO_2_ utilization and *Methanothrix* with acetate consumption. Thus, methanol likely enters the community through a limited number of primary utilizers and is subsequently redistributed to methanogens and dechlorinators through intermediates such as H_2_ and acetate, implying that sustained dechlorination depends on the full syntrophic network rather than on *Dehalobacter* alone (Nelson et al., 2014; Bhatt et al., 2007; Dutta et al., 2022).

Beyond electron-donor transfer, the data are also consistent with corrinoid-dependent support of the dechlorinating guild. Corrinoids (vitamin B_12_-like cofactors) are required for reductive dehalogenase activity in *Dehalobacter* (Wang et al., 2017), and our modeling identified cobalt as an important trace requirement linked to corrinoid and reductive dehalogenase synthesis. Although the present GEMs did not explicitly capture corrinoid exchange, *Acetobacterium sp003260995* and some fermentative community members harbored corrinoid-biosynthetic capacity, making them plausible suppliers of these cofactors to *Dehalobacter* (Men et al., 2015). Together with the coexistence of two *Dehalobacter* populations carrying distinct RDase complements, these results support a model in which chlorinated benzene transformation is carried out by a specialized dechlorinating guild, but depends on surrounding community members for both electron donation and cofactor provision (Alfán-Guzmán et al., 2017; Nelson et al., 2014; Qiao et al., 2019).

### 4.4. Insights from genome-scale modeling

Integrating genome-centric metagenomics with GEM modeling proved valuable for dissecting the metabolic potential and interaction structure of the *Dehalobacter*-containing consortium. This systems-biology framework moves beyond taxonomic description by enabling prediction of functional roles and metabolic interdependencies (Cerk et al., 2024; Carter et al., 2024). In our study, GEM-based analyses identified likely differences in MAG growth under simulated reactor conditions, essential nutrient requirements across major populations, shared versus taxon-specific metabolic capabilities, candidate pairwise ecological interactions, and cross-feeding metabolites such as H_2_, acetate, and CO_2_ that help organize community structure. The models also highlighted shared trace-metal requirements, including cobalt, which is relevant to corrinoid and reductive dehalogenase synthesis, while Jaccard-based comparisons and community flux sampling helped resolve both conserved metabolic structure and heterogeneity in feasible exchange states (Gelbach et al., 2024; Herrmann et al., 2019).

Despite these strengths, GEMs still represent encoded metabolic potential rather than direct *in situ* activity, and therefore do not fully capture regulation, enzyme activity, or environmental heterogeneity (Bernstein et al., 2021). FBA-based approaches also rely on simplifying assumptions, including steady-state behavior and optimization criteria such as biomass maximization or metabolic parsimony, which may not fully reflect biological complexity (Colarusso et al., 2021; Schulz et al., 2021; Orth et al., 2010). Model fidelity is further constrained by annotation quality and gap-filling accuracy, especially for MAG-derived reconstructions (Pan and Reed, 2018; Boer et al., 2024). Although we applied rigorous quality control, including MEMOTE assessment and manual curation of known *Dehalobacter* pathways, some predictions did not fully match experimental observations; for example, some *Bacteroidales* were predicted to grow slowly despite high relative abundance, and the prevalence of parasitic or amensal outcomes contrasted with the long-term stability of the enrichment. Together, these discrepancies suggest that even with flux sampling, static community GEMs likely oversimplify dynamic and spatially structured behavior. Recent developments in dynamic and spatiotemporally resolved metabolic modeling reinforce this point (Moimenta et al., 2025; Pontrelli et al., 2025; Blasche et al., 2021; Li et al., 2022), and future integration of transcriptomic, proteomic, or metabolomic data should further improve the predictive power and biological realism of community-scale models.

### 4.5. Role of methanol as electron donor and carbon source

The use of methanol as the primary electron donor and carbon source strongly influenced the structure and metabolic dynamics of the enrichment cultures. Methanol is a simple C1 substrate that can be metabolized by a restricted subset of anaerobes, including acetogens and some methanogens (Costa and Whitman, 2023; Debabov, 2021). In our metagenomic analysis, *Acetobacterium* was the clearest candidate for primary methanol conversion, whereas *Methanobacterium* and *Methanothrix* were more likely to occupy downstream methanogenic niches. Consistent with this, the reconstructed GEMs supported methanol-associated metabolism in *Acetobacterium*, including the potential conversion of methanol and CO_2_ to acetate via the Wood–Ljungdahl pathway. By contrast, *Methanobacterium* is more plausibly linked to H_2_/CO_2_ utilization, while *Methanothrix* is more plausibly linked to acetate consumption (Thauer et al., 2008). These results suggest that methanol entered the community metabolic network through a limited number of primary utilizers and was subsequently redistributed to dechlorinators and methanogens through intermediates such as H_2_ and acetate.

The flow of electrons from methanol to the terminal electron acceptor (chlorobenzenes via *Dehalobacter*) likely proceeds through intermediate carriers, primarily H_2_ and possibly acetate. While *Dehalobacter* itself is not known to directly utilize methanol, it relies on H_2_ generated by other community members (Nelson et al., 2014). The reliance on methanol therefore requires the presence and activity of organisms capable of converting methanol into H_2_ or other usable electron donors for *Dehalobacter*. This contrasts with communities fed directly with H_2_, lactate, or formate, where the electron donor is more readily available to dechlorinators, potentially leading to different community compositions and interaction networks (Bhatt et al., 2007; Dutta et al., 2022). For example, direct H_2_ feeding might reduce the abundance or necessity of fermentative or acetogenic methanol converters but could increase competition for H_2_ between dechlorinators and methanogens. The use of methanol can select a more complex syntrophic chain, potentially increasing the resilience of the community, but also introducing more potential bottlenecks if methanol conversion becomes limiting. Our modeling results, which show significant roles for *Acetobacterium* and methanogens, support the hypothesis that methanol metabolism is a key structuring factor in this consortium, channeling carbon and electrons to both methanogenesis and dechlorination via interspecies H_2_ and acetate transfer.

### 4.6. Bioremediation implications and engineering consortia

Understanding the metabolic interactions within the *Dehalobacter*-containing consortium has significant implications for optimizing the bioremediation of chlorobenzene-contaminated sites. Incomplete dechlorination of 1,2,4-TCB, often halting at MCB, remains a persistent challenge in anaerobic systems (Nelson et al., 2014; Qiao et al., 2019). Our study, which combines an experimental enrichment capable of dechlorinating MCB to benzene with modeling, highlights how specific community dynamics, such as hydrogen competition, cofactor dependencies, and syntrophic support, can enable or hinder detoxification. For example, managing the availability of electron donors (e.g. methanol dosing or direct H_2_ delivery) may improve reductive dechlorination by alleviating competition and suppressing excessive methanogenesis (Aulenta et al., 2006), while supporting organisms like *Acetobacterium* may be critical as corrinoid producers in settings limited by nutrients (Yan et al., 2013). These insights offer rational targets for enhancing microbial function via biostimulation or bioaugmentation, including the co-introduction of syntrophic partners alongside *Dehalobacter*, a strategy supported by emerging work in the design of synthetic microbial communities (Singha and Shukla, 2023; Jones et al., 2024). Our identification of a core group (*Dehalobacter, Acetobacterium, Methanobacterium, Methanothrix*, and key fermenters) offers a blueprint for such engineered consortia. However, translating laboratory findings into the field must consider site-specific constraints, such as pH, temperature, geochemistry, and cocontaminants, that shape the viability and interactions of microbes (Maphosa et al., 2010; Saiyari et al., 2018). Despite these complexities, the systems-level insights gained here provide a foundational step toward more predictable and resilient bioremediation designs for persistent halogenated pollutants.

In conclusion, this study advances our understanding of the metabolic complexity underlying the anaerobic dechlorination of chlorinated benzenes by a *Dehalobacter*-containing consortium through an integrated systems biology approach. By combining experimental evidence with genome-centric metagenomics and predictive metabolic modeling, we demonstrated MCB-to-benzene conversion, identified coexisting *Dehalobacter* spp., and reconstructed a community-level model that elucidates how microbial interdependencies govern bioremediation outcomes. The findings underscore that effective detoxification is not solely driven by the presence of OHRB, but also by syntrophic networks involving electron donor supply, corrinoid provision, and hydrogen scavenging by methanogens. The coexistence of multiple *Dehalobacter* populations suggests possible niche specialization or redundancy, with implications for the design of robust consortia. Importantly, our application of GEMs provides a predictive framework for exploring metabolic bottlenecks, nutrient dependencies, and the effects of environmental or engineered perturbations. Although further empirical validation is needed, this study exemplifies how system-level approaches can guide the rational design and optimization of microbial communities for the bioremediation of persistent pollutants such as chlorinated benzenes.

## 5. Data and code availability statement

The sequencing data for this study have been deposited in the European Nucleotide Archive (ENA) at EMBL-EBI under accession number `PRJEB112269` (https://www.ebi.ac.uk/ena/browser/view/PRJEB112269). All scripts, final models, and supplementary data used or generated in this study are available at the following GitLab repository: https://git.wur.nl/unlock/projects/dehalobacter-community-modeling.git. Furthermore, the GEMs, as well as the accompanying MEMOTE (Lieven et al., 2020) and FROG Analysis (Raman et al., 2024) reports, can be found in a public repository on BioModels: https://www.ebi.ac.uk/biomodels/, with model identifier: MODEL2605020001.

## 6. Acknowledgments

W.T.S.J., J.J.K., B.N., H.S., P.J.S. and R.K. acknowledge the Dutch Research Council (NWO) and Wageningen University & Research for their financial contribution to UNLOCK (NWO: 184.035.007). The authors thank Dr. Johannes Zimmermann for important suggestions provided during the generation of GEMs. Furthermore, the authors thank Martijn Melissen for his help in creating the phylogenetic tree visualization.

## CRediT authorship contribution statement

**William T. Scott:** Visualization, Validation, Methodology, Investigation, Formal analysis, Conceptualization, Writing – original draft. **Luz A. Puentes Jácome:** Data curation, Methodology, Investigation, Conceptualization, Writing - Proofreading and editing. **Bart Nijsse:** Conceptualization, Formal analysis, Writing - Proofreading and editing. **Jinsong Wang:** Data curation, Writing - Proofreading and editing. **Gerben R. Stouten:** Data curation, Writing - Proofreading and editing. **Jasper J. Koehorst:** Conceptualization, Data curation, Writing - Proofreading and editing. **Hauke Smidt:** Funding acquisition,Supervision, Project administration, Writing - Review and editing. **Elizabeth A. Edwards:** Conceptualization, Project administration, Writing - Review and editing. **Peter J. Schaap:** Conceptualization, Project administration, Writing - Review and editing. **Robbert Kleerebezem:** Conceptualization, Funding acquisition, Project administration, Writing - Review and editing.

